# Absolute proteome quantification in the gas-fermenting acetogen *Clostridium autoethanogenum*

**DOI:** 10.1101/2021.05.11.443690

**Authors:** Kaspar Valgepea, Gert Talbo, Nobuaki Takemori, Ayako Takemori, Christina Ludwig, Alexander P. Mueller, Ryan Tappel, Michael Köpke, Séan Dennis Simpson, Lars Keld Nielsen, Esteban Marcellin

## Abstract

Microbes that can recycle one-carbon (C1) greenhouse gases into fuels and chemicals are vital for the biosustainability of future industries. Acetogens are the most efficient known microbes for fixing carbon oxides CO_2_ and CO. Understanding proteome allocation is important for metabolic engineering as it dictates metabolic fitness. Here, we use absolute proteomics to quantify intracellular concentrations for >1,000 proteins in the model-acetogen *Clostridium autoethanogenum* grown on three gas mixtures. We detect prioritisation of proteome allocation for C1 fixation and significant expression of proteins involved in the production of acetate and ethanol as well as proteins with unclear functions. The data also revealed which isoenzymes are important. Integration of proteomic and metabolic flux data demonstrated that enzymes catalyse high fluxes with high concentrations and high in vivo catalytic rates. We show that flux adjustments were dominantly accompanied with changing enzyme catalytic rates rather than concentrations. Our work serves as a reference dataset and advances systems-level understanding and engineering of acetogens.

## INTRODUCTION

Increasing concerns about irreversible climate change are accelerating the shift to renewable, carbon-free energy production (e.g., solar, wind, fuel cells). However, many fuels and chemicals will stay carbon-based, and thus, technologies for their production using sustainable and renewable feedstocks are needed to transition towards a circular bioeconomy. Moreover, the rising amount of solid waste produced by human activities (e.g., municipal solid waste, lignocellulosic waste) will further endanger our ecosystems’ already critical state. Both challenges can be tackled by using organisms capable of recycling gaseous one-carbon (C1) waste feedstocks (e.g., industrial waste gases [CO_2_, CO, CH_4_], syngas from gasified biomass or municipal solid waste [CO, H_2_, CO_2_]) into fuels and chemicals at industrial scale^1–3^.

As we transition into a new bioeconomy, a key feature of global biosustainability will be the capacity to convert carbon oxides into products at industrial scale. Acetogens are the ideal biocatalysts for this as they use the most energy-efficient pathway, the Wood-Ljungdahl pathway (WLP)^4,5^, for fixing CO_2_ into the central metabolite acetyl-CoA^6–9^ and accept gas (CO, H_2_, CO_2_) as their sole carbon and energy source^5^. Indeed, the model-acetogen *Clostridium autoethanogenum* is already being used as a cell factory in industrial-scale gas fermentation^3,10^. The WLP is considered the first biochemical pathway on Earth^7,11–13^ and continues to play a critical role in the biogeochemical carbon cycle by fixing an estimated 20% of the global CO_2_^6,14^. While biochemical details of the WLP are well described^4,6,15^, a quantitative understanding of acetogen metabolism is just emerging^16,17^. Notably, recent systems-level analyses of acetogen metabolism have revealed mechanisms behind metabolic shifts^18–21^, transcriptional architectures^22,23^, and features of translational regulation^24,25^. However, we still lack an understanding of acetogen proteome allocation through the quantification of proteome-wide intracellular protein concentrations. This fundamental knowledge is required for advancing rational metabolic engineering of acetogen cell factories and for accurate in silico reconstruction of their phenotypes using metabolic models^1,2^.

Quantitative description of an organism’s proteome allocation through absolute proteome quantification is valuable in several ways. Firstly, it enables us to understand prioritisation of the energetically costly proteome resources among functional protein categories, metabolic pathways, and single proteins^26,27^. This may also identify relevant proteins with unclear functions and high abundances. Secondly, some metabolic fluxes can be catalysed by isoenzymes and a comparison of their intracellular concentrations can indicate which are likely relevant in vivo and are thus targets for genetic perturbation experiments to validate in vivo functionalities^28^. Thirdly, integration of absolute proteomics and metabolic flux data enable the estimation of apparent in vivo catalytic rates of enzymes (k_app_)^26,29^, which can be used to identify less-efficient enzymes as targets for improving pathways through metabolic and protein engineering. Absolute proteomics data also contribute to the curation of accurate genome-scale metabolic models.

Absolute proteome quantification is generally performed using label-free mass-spectrometry (MS) approaches without spike-in standards^30,31^. The major limitation of this approach is that accuracy of label-free estimated protein concentrations cannot be determined. Furthermore, the optimal model to convert MS signals (e.g., spectral counts, peak intensities) into protein concentrations remains unknown^32–34^. Label-based approaches using stable-isotope labelled (SIL) spike-ins of endogenous proteins are thus preferred for reliable absolute proteome quantification. This strategy relies on accurate absolute quantification of a limited set of intracellular proteins (i.e., anchors) using SIL spike-ins to establish a linear correlation between protein concentrations and their measured MS intensities^32^. Studies with the latter approach have determined a 1.5 □ 2.4-fold error for label-free estimation of proteome-wide protein concentrations in multiple organisms^28,35–41^.

The aim of our work was to perform reliable absolute proteome quantification for the first time in an acetogen. We employed a label-based MS approach using SIL-protein spike-in standards to quantify SIL-based concentrations for 16 key proteins and label-free-based concentrations for >1,000 *C. autoethanogenum* proteins during autotrophic growth on three gas mixtures. This allowed us to explore global proteome allocation, uncover isoenzyme usage in central metabolism, and quantify regulatory principles associated with estimated k_app_s. Our work provides an important reference dataset and advances the systems-level understanding and engineering of the ancient metabolism of acetogens.

## RESULTS

### Absolute proteome quantification framework in the model-acetogen *C. autoethanogenum*

We performed absolute proteome quantification from autotrophic steady-state chemostat cultures of *C. autoethanogenum* grown on three different gas mixtures: CO, syngas (CO+CO_2_+H_2_), or CO+H_2_ (termed “high-H_2_ CO”) described before^18,19^. Briefly, four biological cultures of each gas mixture were grown anaerobically on a chemically defined medium at 37 °C, pH of 5, and dilution rate ∼1 day^-1^ (specific growth rate ∼0.04 h^-1^) without the use of heavy SIL substrates. The absolute proteome quantification framework (Fig. 1) was built on using 19 synthetic heavy SIL-variants of key *C. autoethanogenum* proteins covering central metabolism (Supplementary Table 1). The SIL-protein standards were spiked in for quantification of intracellular concentrations of their endogenous light counterparts. This framework ensures accurate absolute quantification compared to commonly used peptide spike-ins. Spiking cell lysates with protein standards before sample clean-up and protein digestion accounts for errors accompanying these critical steps^30,31,42,43^. Furthermore, selection of peptides ensuring accurate absolute protein quantification without prior MS data is challenging as its difficult to predict which peptides “fly” well^30,31,42,43^. In contrast, all proteotypic peptides from a protein spike-in can be used for quantification.

**Fig. 1.**
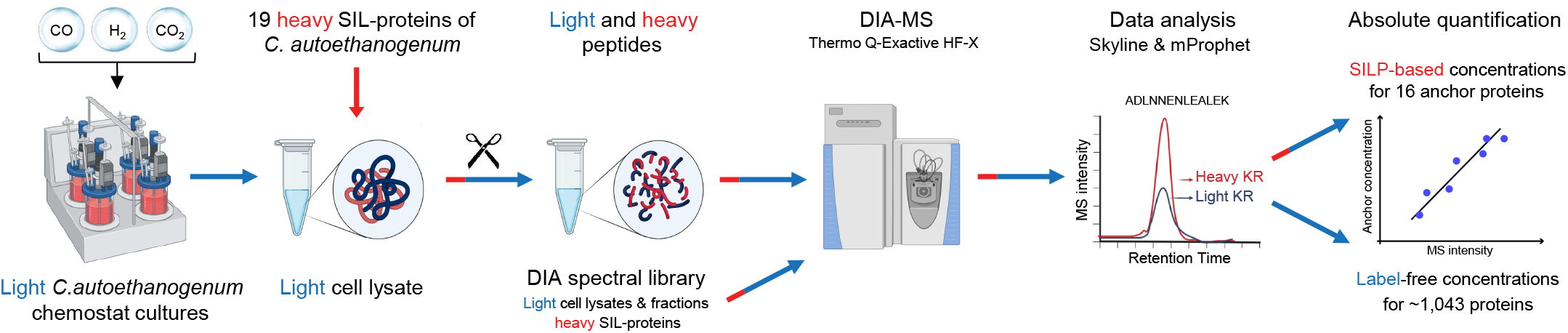
Absolute proteome quantification framework in *C. autoethanogenum*. Absolute proteome quantification in light (no stable-isotope labelled [SIL] substrates) autotrophic *C. autoethanogenum* chemostat cultures was built on using 19 synthetic heavy SIL-protein spike-in standards and data-independent acquisition (DIA) mass spectrometry (MS) analysis. Culture samples with SIL-protein spike-ins and samples for DIA spectral library were analysed by DIA MS. Subsequent stringent data analysis allowed to quantify intracellular concentrations for 16 key *C. autoethanogenum* proteins using light-to-heavy ratios between endogenous and spike-in DIA MS intensities. These 16 key proteins were further used as anchor proteins for label-free estimation of ∼1,043 protein concentrations through establishing a linear correlation between protein concentrations and their measured MS intensities. Some parts created with BioRender.com.

We synthesised heavy-labelled lysine and arginine SIL-proteins using a cell-free wheat germ extract platform as described previously^18,44,45^ and quantified standard stocks using parallel reaction monitoring (PRM) MS. Next, proteins were extracted from culture samples using an optimised protocol maximising extraction yield^18^ followed by spike-in of the 19 heavy SIL-proteins into light cell lysates. We then used a data-independent acquisition (DIA) MS approach^46^ to quantitate 1,243 proteins of *C. autoethanogenum* across 12 samples (quadruplicate cultures of three gas mixtures) using a comprehensive spectral library consisting of whole-cell lysates, lysate fractions, and spike-in SIL-proteins. Finally, we quantified intracellular concentrations for 16 key *C. autoethanogenum* proteins using light-to-heavy ratios between endogenous and spike-in DIA MS intensities and further used these 16 as anchor proteins for label-free estimation of ∼1,043 protein concentrations through establishing a linear correlation between protein concentrations and their measured MS intensities^32^. We express intracellular protein concentrations in nanomoles of protein per gram of dry cell weight (nmol/gDCW).

### Absolute quantification of 16 anchor protein concentrations

To ensure high confidence absolute quantification of anchor protein concentrations from the DIA MS data, we employed stringent criteria on top of the automated mProphet peak picking algorithm^47^ within the software Skyline^48^. We also performed a dilution series experiment for each SIL-protein to increase accuracy (see Methods for details). Briefly, we kept only peaks with Gaussian shapes and without interference and precursors with highest Skyline quality metrics. Importantly, only peptides whose signal were above the lower limit of quantification (LLOQ) and within the linear dynamic quantification range in the dilution series experiment were used for anchor protein quantification (Supplementary Table 2). We thus used 106 high-confidence peptides for the absolute quantification of 16 anchor protein concentrations (Table 1; see also Fig. 5). High confidence of the intracellular concentrations for these key *C. autoethanogenum* proteins of central metabolism is supported both by the low average 11% coefficient of variation (CV) between biological quadruplicate cultures (Table 1) and the average 22% CV between different peptides of single proteins (Supplementary Table S2).

**Fig. 5.**
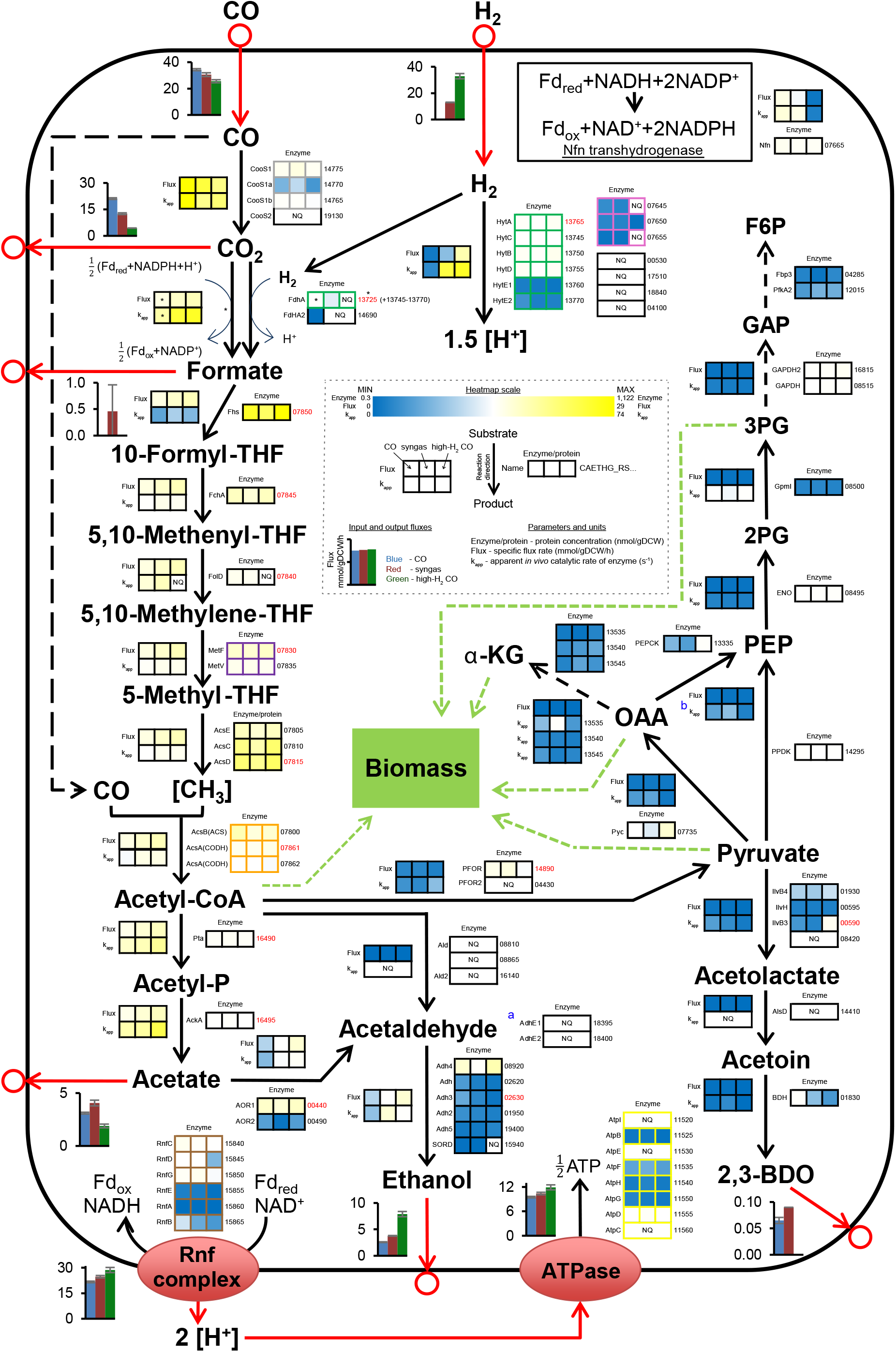
Quantitative systems-level view of acetogen central metabolism. Enzyme concentrations (nmol/gDCW), apparent in vivo catalytic rates of enzymes (k_app_; s^-1^), and metabolic flux rates (mmol/gDCW/h) are shown for *C. autoethanogenum* steady-state chemostat cultures grown on three gas mixtures. See dashed inset for bar chart and heatmap details. Enzyme concentration and k_app_ data are average of biological replicates. Proteins forming a complex are highlighted with non-black borders (FdhA forms a complex with HytA–E for direct CO_2_ reduction with H_2_; CooS1 is expected to form a complex with CooS1a and b as they are encoded from the same operon). For reactions with isoenzymes, k_app_ is for the enzyme with the highest concentration ranking (top location on enzyme heatmap), see Methods for details. Flux data from ref.^18^ are average of biological replicates and error bars denote standard deviation. Arrows show direction of calculated fluxes; red arrow denotes uptake or secretion. Gene/protein IDs right of enzyme concentration heatmaps are preceded with CAETHG_RS and red font denotes concentrations determined using stable-isotope labelled (SIL) protein spike-in standards (i.e., anchor proteins). Asterisk denotes data for redox-consuming CO_2_ reduction to formate solely by FdhA without the use of H_2_ during growth on CO. ^a^Bifunctional acetaldehyde/alcohol dehydrogenase (acetyl-CoA→ethanol); ^b^Flux into PEP from OAA and pyruvate is merged and k_app_ is for PEPCK. See Supplementary Table S3 for gene/protein ID, proposed name, description, and label-free protein concentrations. See Table 1 for anchor protein concentrations. See Supplementary Table S5 for k_app_ and flux data. See ref.^18^ for cofactors of reactions and metabolite abbreviations. *gDCW*, gram of dry cell weight, *NQ*, not quantified.

### Label-free estimation of proteome-wide protein concentrations

Both high quality proteomics data and suitable anchor proteins are required for reliable label-free absolute proteome quantification. Our proteome-wide DIA MS data were highly reproducible with an average Pearson correlation coefficient of R = 0.99 between biological replicates (Fig. 2a and Supplementary Fig. 1). We also found our anchor proteins suitable as their concentrations spanned across three orders of magnitude, and the summed mass accounted for ∼⅓ of the peptide mass injected into the mass spectrometer (Table 1 and Fig. 2b). We used the 16 anchor proteins (with 106 peptides) to determine the optimal label-free quantification model with the best linear fit between anchor protein concentrations and their measured DIA MS intensities using the aLFQ R package^49^ as described before for SWATH MS^28^ (Fig. 2c). Notably, we detected an average 1.5-fold cross-validated mean fold-error (CV-MFE; bootstrapping) for the label-free estimated anchor protein concentrations across samples (Fig. 2d). The errors were distributed normally (Supplementary Fig. 2) with an average 95% CI of 0.3 (Fig. 2d). We then applied the optimal label-free quantification model to estimate ∼1,043 protein concentrations in *C. autoethanogenum* (Supplementary Table 3).

**Fig. 2.**
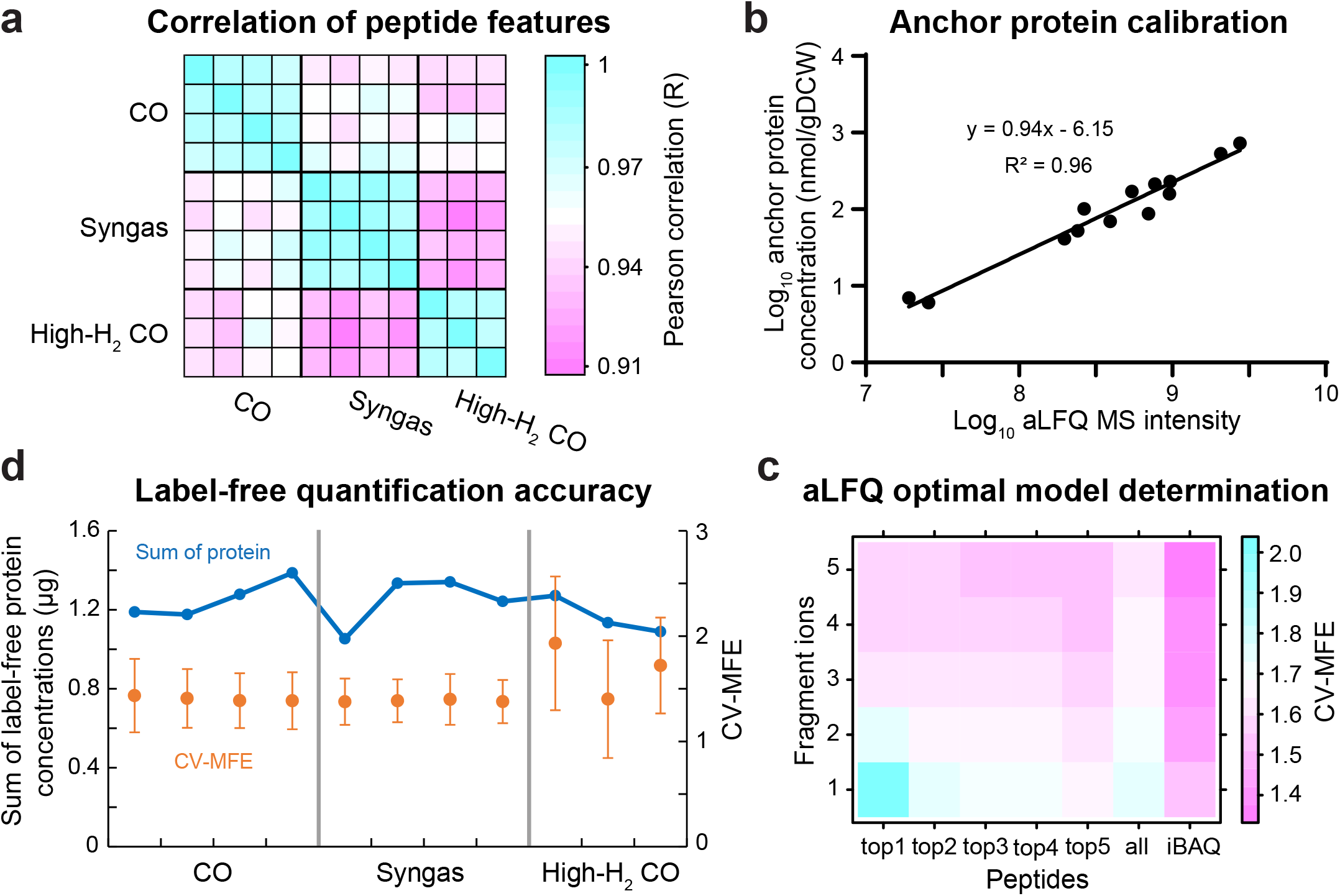
Label-free estimation of proteome-wide protein concentrations. **a** Correlation of peptide mass spectrometry (MS) feature intensities between biological replicate cultures of the three gas mixtures. **b** Linear correlation between anchor protein concentrations and their measured MS intensities for one syngas culture. *gDCW*, gram of dry cell weight, *aLFQ*, absolute label-free quantification. **c** Errors of different label-free quantification models for the linear fit between anchor protein concentrations and their measured MS intensities determined by bootstrapping using the aLFQ R package^49^ for one syngas culture. *CV-MFE*, cross-validated mean fold-error. **d** Label-free quantification error of optimal model (orange) and total proteome mass (blue) across samples. Error bars denote 95% CI.

Prior to the detailed analysis of proteome-wide protein concentrations, we further evaluated our label-free data accuracy beyond the 1.5-fold CV-MFE determined above. Firstly, the total proteome mass (1.2±0.1 μg; average ± standard deviation) closely matched the 1 μg peptide mass injected into the mass spectrometer (Fig. 2d). The data were also supported by a strong correlation between estimated protein concentrations and expected stoichiometries for equimolar (Fig. 3) and non-equimolar protein complexes (Supplementary Fig. 3). Notably, absolute protein concentrations of syngas cultures correlated well (R = 0.65) with their respective absolute transcript expression levels determined before^19^ (Supplementary Fig. 4). This result is similar to the correlations of absolute data seen in other steady-state cultures^26,50^. Altogether, we present the first absolute quantitative proteome dataset for a gas-fermenting acetogen that includes SIL-based concentrations for 16 key proteins and label-free estimates for over 1,000 *C. autoethanogenum* proteins during growth on three gas mixtures.

**Fig. 3.**
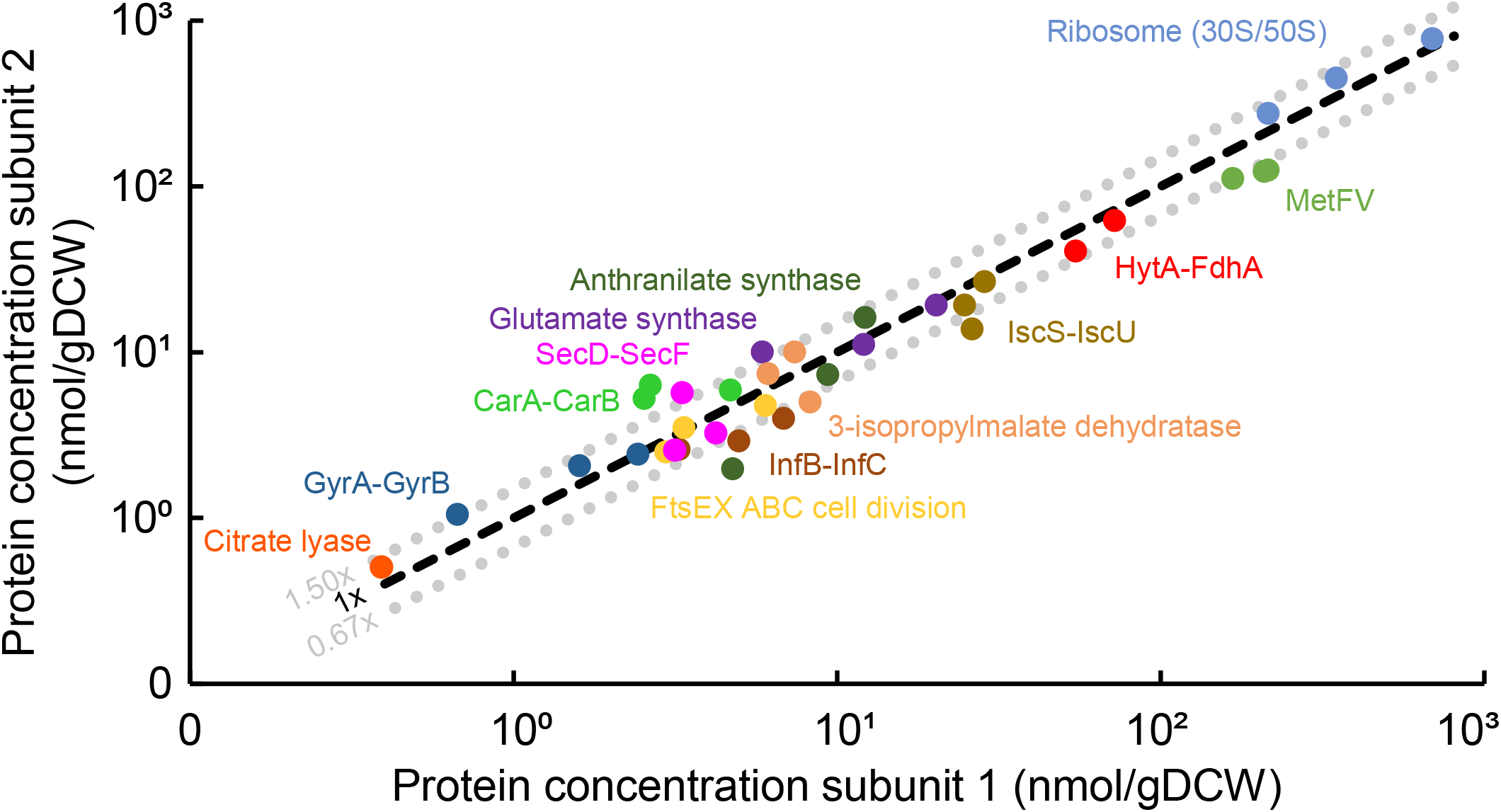
Strong correlation between protein concentrations and expected stoichiometries for equimolar protein complexes. Grey dotted lines denote the average 1.5-fold cross-validated mean fold-error (CV-MFE) of label-free protein concentrations. Label-free protein concentrations are plotted, except for the HytA-FdhA complex, which was quantified using stable-isotope labelled protein spike-ins. Data points of the same colour represent gas mixtures. See Methods for details on expected protein complex stoichiometries. See Supplementary Table S3 for gene/protein ID, proposed name, description, and label-free data. See Table 1 for HytA-FdhA data. *gDCW*, gram of dry cell weight.

### C1 fixation dominates global proteome allocation

Global proteome allocation amongst functional gene classifications was explored using proteomaps^27^ and KEGG Orthology identifiers (KO IDs)^51^. The “treemap” structure defining the four-level hierarchy of our proteomaps (Supplementary Table 4) also included manually curated categories to accurately reflect acetogen metabolism (e.g., C1 fixation/WLP, Hydrogenases). As expected for autotrophic growth of an acetogen, the C1 fixation (Fig. 4) or WLP (Supplementary Fig. 5) categories dominated the proteome allocation with a ∼⅓ fraction, compared to Carbohydrate metabolism or Glycolysis/Gluconeogenesis. Notably, the data show that two genes–dihydrolipoamide dehydrogenase (LpdA; CAETHG_RS07825) and glycine cleavage system H protein (GcvH; RS07795)–encoded by the WLP gene cluster were translated at very high levels (Fig. 4). This is important as both have unknown functions in *C. autoethanogenum* metabolism. Significant investment in expression of proteins involved in acetate and ethanol production (Supplementary Fig. 5) is consistent with ⅓-to-⅔ of fixed carbon channelled into these two growth by-products across the three gas mixtures^18,19^. The 11% proteome fraction of category Translation (Fig. 4) is expected for cells growing at a specific growth rate ∼0.04 h^-1^ based on absolute proteomics data from *Escherichia coli*^26,39,52^. The notable proteome allocation for Amino acid metabolism and particularly the high abundance of ketol-acid reductoisomerase (IlvC; RS00580) are surprising since metabolic fluxes through 2,3-butanediol and branched-chain amino acid pathways were low under these growth conditions^18,19^. In addition, numerous proteins with unknown or unclear functions (coloured grey in proteomaps) are highly expressed (e.g., RS12590, RS08610, RS08145), highlighting the need for global mapping of genotype-phenotype relationships in acetogens. In general, proteome allocation was highly similar between the three gas mixtures (Supplementary Table 3). This result is unsurprising given the few relative protein expression differences detected previously^18^.

**Fig. 4.**
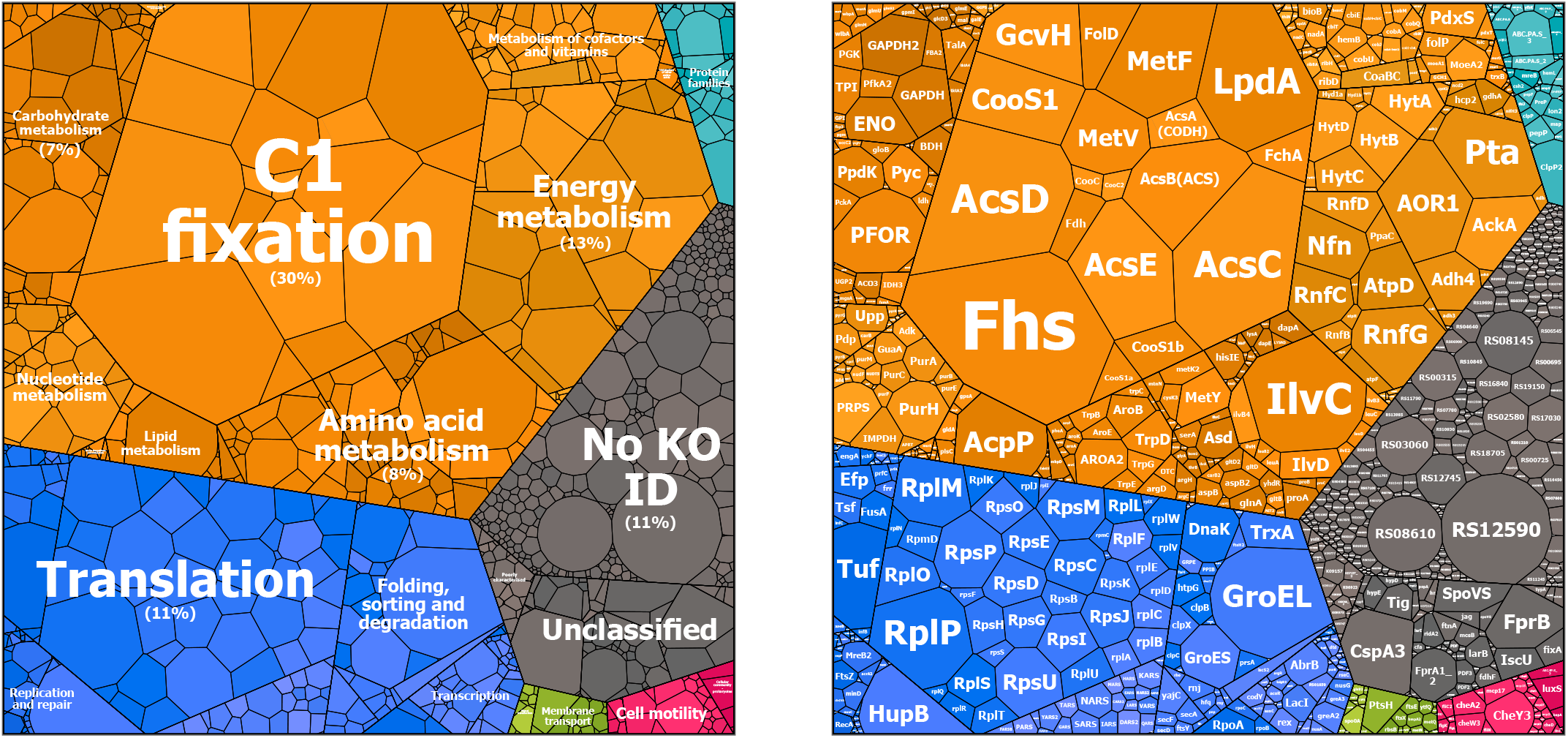
Proteomaps uncover global proteome allocation. Left proteomap shows proteome allocation amongst functional gene classification categories (KEGG Orthology identifiers [KO IDs] ^51^) at level two of the four-level “treemap” hierarchy structure (Supplementary Table S4). Right proteomap shows proteome allocation at the level of single proteins (level four of “treemap”). See Supplementary Fig. S5 for proteomaps of levels one and three of “treemap”. Area of the tile is proportional to protein concentration. Colours denote level one categories of “treemap”. Proteomaps visualise average concentrations of syngas cultures while category percentages are average of three gas mixtures (shown for categories with a fraction >5%). See Supplementary Table S3 for gene/protein ID, proposed name, description, and label-free protein concentrations.

### Enzyme usage revealed in central metabolism

Next, we focused on uncovering enzyme usage in acetogen central metabolism (Fig. 5). This contains enzymes of the WLP, acetate, ethanol, and 2,3-butanediol production pathways, hydrogenases, and the Nfn transhydrogenase, which together carry >90% of the carbon and most of the redox flow in *C. autoethanogenum*^18–20^. Multiple metabolic fluxes in these pathways can be catalysed by isoenzymes and absolute proteomics data can indicate which of the isoenzymes are likely relevant in vivo. While the carbon monoxide dehydrogenase (CODH) AcsA (RS07861□62) that forms the bifunctional CODH/ACS complex with the acetyl-CoA synthase^53^ (AcsB; RS07800) is essential for *C. autoethanogenum* growth on gas as confirmed in mutagenesis studies^54^, the higher concentrations of the dispensable monofunctional CODH CooS1 (RS14775) suggest it may also play a role in CO oxidation (Fig. 4, 5), in addition to CO_2_ reduction^54^. Additionally, our proteomics data show high abundance of the primary acetaldehyde:ferredoxin oxidoreductase (AOR1; RS00440) and this support the emerging understanding that in *C. autoethanogenum* ethanol is dominantly produced using the AOR1 activity via acetate, instead of directly from acetyl-CoA via acetaldehyde using mono- or bifunctional activities^18,19,55,56^ (Fig. 5). Furthermore, the data suggest that the specific alcohol dehydrogenase (Adh4; RS08920) is responsible for reducing acetaldehyde to ethanol, a key reaction in terms of carbon and redox metabolism. The high abundance of the electron-bifurcating hydrogenase HytA-E complex (RS13745□70) compared to alternative hydrogenases confirms that it is the main H_2_-oxidiser^57,58^ (Fig. 5). This is consistent with the fact that in the presence of H_2_ all the CO_2_ fixed by the WLP is reduced to formate using H_2_ by the HytA-E and formate dehydrogenase (FdhA; RS13725) enzyme complex activity^18,19^. Despite the proteomics evidence, genetic perturbations are required to determine condition-specific in vivo functionalities of isoenzymes in acetogens unequivocally.

The overall most abundant protein was the formate-tetrahydrofolate ligase (Fhs; RS07850), a key enzyme in the WLP (Fig. 5, 4). Despite the high abundance, its expression might still be rate-limiting (see below). Another key enzyme for acetogens is AcsB because of its essentiality for acetyl-CoA synthesis by the CODH/ACS complex. AcsB is linked to the WLP by the corrinoid iron sulfur proteins AcsC (RS07810) and AcsD (RS07815) that supply the methyl group to AcsB. Interestingly, the ratio of AcsCD-to-AcsB increased from 1.7 (CO) to 2.3 (syngas) to 2.9 (high-H_2_ CO), suggesting that the primary role of the CODH/ACS complex shifted from CO oxidation towards acetyl-CoA synthesis, likely because increased H_2_ uptake could replace the supply of reduced ferredoxin from CO oxidation. Concurrently, the Nfn transhydrogenase (RS07665) levels that act as a redox valve in acetogens^20^ are maintained high (Fig. 5), potentially to rapidly respond to redox perturbations. We conclude that absolute quantitative proteomics can significantly contribute to a systems-level understanding of metabolism, particularly in less-studied organisms.

### Integration of absolute proteomics and flux data yields in vivo enzyme catalytic rates

Absolute proteomics data enable estimation of intracellular catalytic working rates of enzymes when metabolic flux rates are known^26,29^. We thus calculated apparent in vivo catalytic rates of enzymes, denoted as k_app_ (s^-1^)^26^, as the ratio of specific flux rate (mmol/gDCW/h) determined before^18^ and protein concentration (nmol/gDCW) (see Methods). This produced k_app_ values for 13 and 48 enzymes/complexes using either anchor or label-free protein concentrations, respectively (Supplementary Table 5 and Fig. 5, 6). The first two critical steps for carbon fixation in the methyl branch of the WLP (i.e., CO to formate) are catalysed at high rates (Fig. 5). Notably, FdhA showed a k_app_∼30 s^-1^ for CO_2_ reduction without H_2_ during growth on CO only, which is similar to in vitro k_cat_ data of formate dehydrogenases in other acetogens^59,60^. Interestingly, the next step of formate reduction was catalysed potentially by a less-efficient enzyme–Fhs–as its k_app_ of ∼3 s^-1^ is significantly lower compared to other WLP enzymes (Fig. 5). Overall, enzymes catalysing reactions in high flux pathways such as the WLP and acetate and ethanol production have higher k_app_s than those of downstream from conversion of acetyl-CoA to pyruvate (Fig. 5). Indeed, enzymes catalysing high metabolic fluxes in *C. autoethanogenum* have both higher concentrations and higher catalytic rates compared to enzymes catalysing lower fluxes as both specific flux rates and enzyme concentrations (Kendall’s τ = 0.56, p-value = 5×10^−9^) and flux and k_app_ (τ = 0.45, p-value = 2×10^−6^) were significantly correlated (Fig. 6a), as seen before for other organisms^26,61^.

**Fig. 6.**
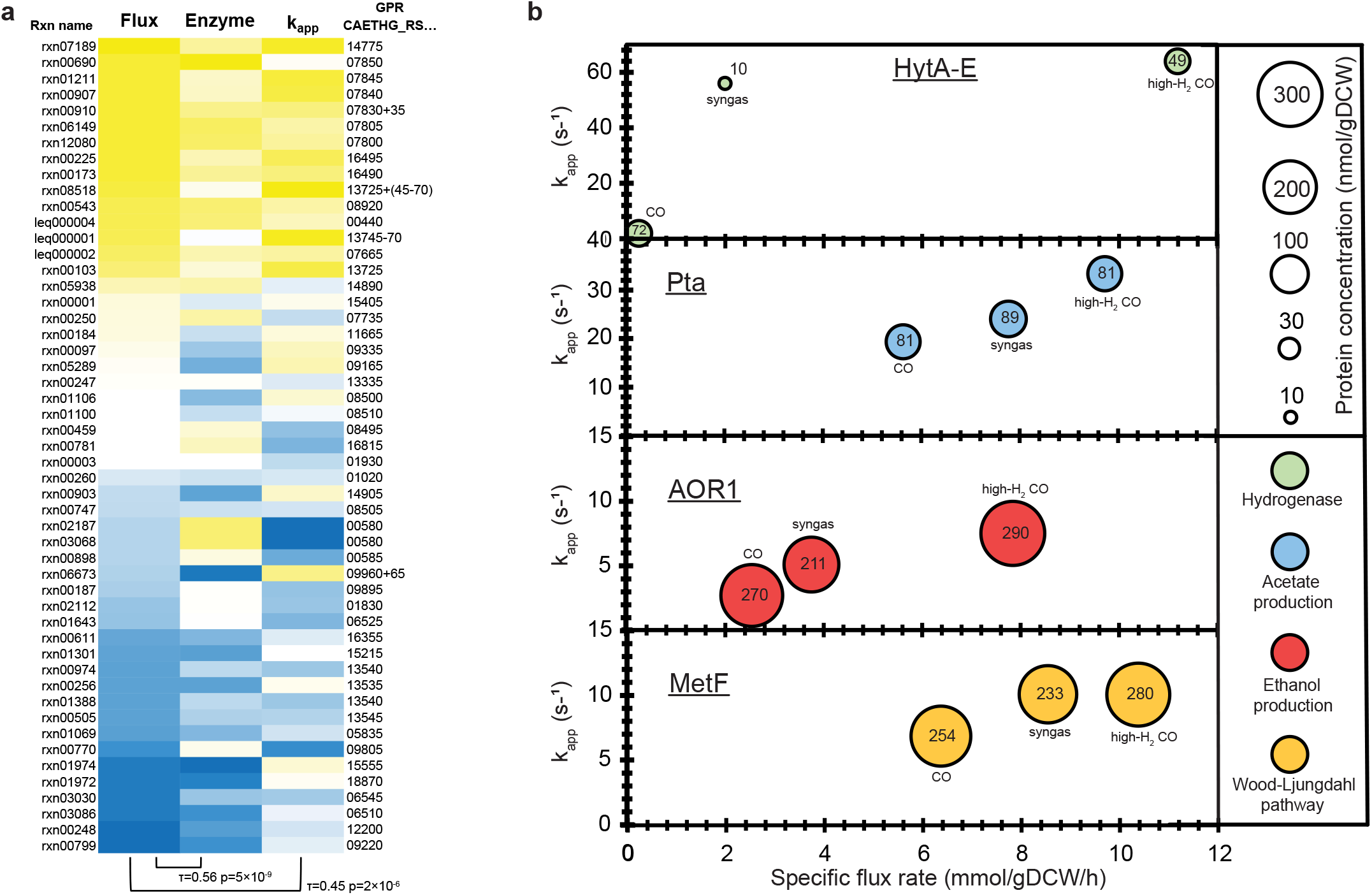
Regulatory principles of apparent in vivo catalytic rates of enzymes (k_app_) and metabolic flux throughput. **a** Enzymes catalysing higher metabolic flux rates have both higher concentrations and higher k_app_s. Yellow and blue denote high and low values, respectively. Kendall’s τ correlations with significance p-values between respective pairs are shown below heatmap. See Supplementary Table S5 for flux rate, enzyme concentration, and k_app_ data, and for description of reaction names (Rxn name) and gene-protein-reaction (GPR) associations. **b** Control of metabolic flux throughput through k_app_ changes for high flux pathways. See also Fig. 5. *gDCW*, gram of dry cell weight.

Having acquired absolute proteomics data for *C. autoethanogenum* growth on three gas mixtures with different metabolic flux profiles also allowed us to determine the impact of change in enzyme concentration and its catalytic rate for adjusting metabolic flux rates. Two extreme examples are the reactions catalysed by the HytA-E (Fig. 6b) and the Nfn (Fig. 5) complexes where flux adjustments were accompanied with large changes in k_app_s rather than in enzyme concentrations. Flux changes in high flux pathways such as the WLP and acetate and ethanol production also coincided mainly with k_app_ changes (Fig. 6b). This principle seems to be dominant in *C. autoethanogenum* as 90% of flux changes were not regulated through enzyme concentrations (i.e., post-translational regulation; Supplementary Table 6) when comparing all statistically significant flux changes between the three gas mixtures with respective enzyme expression changes (see Methods).

## DISCUSSION

The looming danger of irreversible climate change and harmful effects of solid waste accumulation are pushing humanity to develop and adopt sustainable technologies for renewable production of fuels and chemicals and for waste recycling. Acetogen gas fermentation offers great potential to tackle both challenges through recycling waste feedstocks (e.g., industrial waste gases, gasified biomass or municipal solid waste) into fuels and chemicals^1,2^. Although the quantitative understanding of acetogen metabolism has recently improved^16,17^, a quantitative description of acetogen proteome allocation was missing. This is needed to advance their metabolic engineering into superior cell factories and accurate in silico reconstruction of their phenotypes^1,2^. Thus, we performed absolute proteome quantification in the model-acetogen *C. autoethanogenum* grown autotrophically on three gas mixtures.

Our absolute proteome quantification framework relied on SIL-protein spike-in standards and DIA MS analysis to ensure high confidence of the determined intracellular concentrations for 16 key *C. autoethanogenum* proteins. We further used these proteins as anchor proteins for label-free estimation of >1,000 protein concentrations. This enabled us to determine the optimal label-free quantification model for our data to infer protein concentrations from MS intensities, which remains unknown in common label-free approaches not utilising spike-in standards^32,33^. More importantly, label-free estimated protein concentrations using the latter approach are questionable as their accuracy cannot be determined. We determined an excellent average error of 1.5-fold for our label-free estimated protein concentrations based on 16 anchor proteins and a bootstrapping approach. This error is in the same range as described in previous studies using SIL spike-in standards for absolute proteome quantification^28,35–41^. Further, we also observed a good match both between estimated and injected proteome mass into the mass spectrometer and between protein concentrations and expected protein complex stoichiometries. We conclude that label-free estimation of proteome-wide protein concentrations using SIL-protein spike-ins and state-of-the-art MS analysis is reasonably accurate.

Quantification of acetogen proteome allocation during autotrophic growth expectedly showed prioritisation of proteome resources for fixing carbon through the WLP, in line with transcript expression data in *C. autoethanogenum*^19,56^. The allocation of one third of the total proteome for C1 fixation is higher than proteome allocation for carbon fixation through glycolysis during heterotrophic growth of other microorganisms^26,36^. High abundances of other key enzymes of acetogen central metabolism were also expected as the WLP, acetate and ethanol production pathways, hydrogenases, and the Nfn transhydrogenase carry >90% of the carbon and most of the redox flow in *C. autoethanogenum*^18–20^. However, very high expression of the two genes–LpdA and GcvH–of the WLP gene cluster with unknown functions in *C. autoethanogenum* is striking, raising the question whether their function in *C. autoethanogenum* could also be to link WLP and glycine synthase-reductase pathways, as recently proposed for another acetogen^62^. Since many other proteins with unknown or unclear functions were also highly abundant, global mapping of genotype-phenotype relationships in acetogens is much needed.

The in vivo functionalities of isoenzymes are not clear for multiple key metabolic fluxes in acetogen central metabolism and absolute proteomics data can indicate which isoenzymes are likely relevant. Oxidation of CO or reduction of CO_2_ is a fundamental step for all acetogens and known to be catalysed by three CODHs in *C. autoethanogenum*^54^. Though only AcsA that forms the bifunctional CODH/ACS complex with the acetyl-CoA synthase^53^ is essential for growth on gas^54^, we detected higher concentrations of the monofunctional CODH CooS1, which deletion strain shows intriguing phenotypes^54^. Concurrently, our data suggest that prioritisation of CODH/ACS activity between CO oxidation and acetyl-CoA synthesis is sensitive to H_2_ availability. Thus, further studies are required to decipher condition-dependent functionalities of CODHs. In addition to CODHs, the biochemical understanding of ethanol production is important in terms of both carbon and redox metabolism. Our data confirm that in *C. autoethanogenum* ethanol is predominantly produced via acetate by AOR1^18,19,55,56^ and more importantly, indicate for the first time that AOR1 activity is followed by Adh4 (previously characterised as butanol dehydrogenase^63^) for reduction of acetaldehyde to ethanol. These observations call for large-scale genetic perturbation experiments to determine unequivocally the condition-specific in vivo functionalities of isoenzymes in acetogens.

Absolute proteomics data offer a unique opportunity to estimate apparent in vivo catalytic rates of enzymes (k_app_)^26,29^ if also metabolic flux data are available. These data are particularly valuable for more accurate in silico reconstruction of phenotypes using protein-constrained genome-scale metabolic models^64,65^. While in vitro k_cat_ and in vivo k_app_ data generally correlate^29^, models using maximal k_app_ values show better prediction of protein abundances^66^. Furthermore, information of k_app_s can infer less-efficient enzymes as targets for improving pathways through metabolic and protein engineering. For example, protein engineering of Fhs (catalysing formate reduction) might improve WLP throughput and carbon fixation since its k_app_ was significantly lower compared to other pathway enzymes. At the same time, the large change in k_app_s of the abundant electron-bifurcating hydrogenase HytA-E and the Nfn transhydrogenase complexes indicate capacity for the cells to rapidly respond to H_2_ availability and redox perturbations, which may be critical for metabolic robustness of acetogens^20^. Overall, we detected both higher concentrations and k_app_s for enzymes catalysing higher metabolic fluxes, which is believed to arise from an evolutionary push towards reducing protein production costs for enzymes carrying high flux^61^. The observation that 90% of flux changes in *C. autoethanogenum* were not regulated through changes in enzyme concentrations is not surprising for a metabolism that operates at the thermodynamic edge of feasibility^16,17^ since post-translational regulation of fluxes is energetically least costly. Further research is needed to identify which mechanism from post-translational protein modification, allosteric regulation, or substrate concentration change is responsible for post-translational regulation of fluxes.

We have produced the first absolute proteome quantification in an acetogen and thus provided understanding of global proteome allocation, isoenzyme usage in central metabolism, and regulatory principles of in vivo enzyme catalytic rates. This fundamental knowledge has potential to advance both rational metabolic engineering of acetogen cell factories and accurate in silico reconstruction of their phenotypes^1,2,64,65^. Our study also highlights the need for large-scale mapping of genotype-phenotype relationships in acetogens to infer in vivo functionalities of isoenzymes and proteins with unknown or unclear functions. This absolute proteomics dataset serves as a reference towards a better systems-level understanding of the ancient metabolism of acetogens.

## METHODS

### Bacterial strain and culture growth conditions

Absolute proteome quantification was performed from high biomass concentration (∼1.4 gDCW/L) steady-state autotrophic chemostat cultures of *C. autoethanogenum* growing on three different gas mixtures with culturing conditions described in our previous works^18,19^. Briefly, four biological replicate chemostat cultures of *C. autoethanogenum* strain DSM 19630 were grown on a chemically defined medium (without yeast extract) either on CO (∼60% CO and 40% Ar), syngas (∼50% CO, 20% H_2_, 20% CO_2_, and 10% N_2_/Ar), or CO+H_2_, termed “high-H_2_ CO”, (∼15% CO, 45% H_2_, and 40% Ar) under strictly anaerobic conditions. The bioreactors were maintained at 37 °C, pH of 5, and dilution rate ∼1 day^-1^ (specific growth rate ∼0.04 h^-1^).

### Cell-free synthesis of stable-isotope labelled protein standards

Twenty proteins covering *C. autoethanogenum* central carbon metabolism, the HytA-E hydrogenase, and a ribosomal protein (Supplementary Table 1) were selected for cell-free synthesis of SIL-proteins as described in ref.^18^. Briefly, genes encoding for these proteins were synthesised by commercial gene synthesis services (Biomatik). Target genes were sub-cloned into the cell-free expression vector pEUE01-His-N2 (CellFree Sciences) and transformed into *Escherichia coli* DH5α from which plasmid DNA was extracted and purified. Correct gene insertion into the pEUE01-His-N2 was verified by DNA sequencing. Subsequently, cell-free synthesis of His-tag fused *C. autoethanogenum* proteins was performed using the bilayer reaction method with the wheat germ extract WEPRO8240H (CellFree Sciences) as described previously^44,45^. mRNAs for cell-free synthesis were prepared by an in vitro transcription reaction while in vitro translation of target proteins was performed using a bilayer reaction where the translation layer was supplemented with L-Arg-^13^C_6_,^15^N_4_ and L-Lys-^13^C_6_,^15^N_2_ (Wako) at final concentrations of 20 mM to achieve high efficiency (>99 %) for stable-isotope labelling of proteins. The in vitro synthesised SIL-protein sequences also contained an N-terminal amino acid sequence GYSFTTTAEK that was later used as a tag for quantification of the SIL-protein stock concentration. Subsequently, SIL-proteins were purified using the Ni-Sepharose High-Performance resin (GE Healthcare Life Sciences) and precipitated using methanol:chloroform:water precipitation in Eppendorf Protein LoBind^®^ tubes. Lastly, precipitated SIL-proteins were reconstituted in 104 μL of 8 M urea ([UA]; Sigma-Aldrich) in 0.1 M Trizma^®^ base (pH 8.5) by vigorous vortexing and stored at -80 °C until further use.

### Absolute quantification of SIL-protein standards using PRM MS

Concentrations of the twenty synthesised SIL-protein standard stocks were determined using PRM MS preceded by in-solution digestion of proteins and sample desalting and preparation for MS analysis.

#### Sample preparation

Only Eppendorf Protein LoBind^®^ tubes and pipette tips were used for all sample preparation steps. Firstly, 20 μL of UA was added to 4 μL of the SIL-protein standard stock used to determine the stock concentration, and the mix was vortexed. Then, 1 μL of 0.2 M DTT (Promega) was added, followed by vortexing and incubation for 1 h at 37 °C to reduce disulphide bonds. Sulfhydryl groups were alkylated with 2 µL of 0.5 M iodoacetamide (IAA; Sigma-Aldrich), vigorous vortexing, and incubation for 30 min at room temperature in the dark. Next, 75 µL of 25 mM ammonium bicarbonate was added to dilute UA down to 2 M concentration. Subsequently, 2 pmol (2 µL of stock) of the non-labelled AQUA^®^ peptide HLEAAKGYSFTTTAEKAAELHK (Sigma-Aldrich) containing the quantification tag sequence GYSFTTTAEK was added to enable quantification of SIL-protein stock concentrations using MS analysis based on the ratio of heavy-to-light GYSFTTTAEK signals (see below). Protein digestion was performed for 16 h at 37 °C with 0.1 µg of Trypsin/Lys-C mix (1 µL of stock; Promega) and stopped by lowering pH to 3 by the addition of 5 µL of 10% (v/v) trifluoroacetic acid (TFA).

Samples were desalted using C_18_ ZipTips (Merck Millipore) as follows: the column was wetted using 0.1% (v/v) formic acid (FA) in 100% acetonitrile (ACN), equilibrated with 0.1% FA in 70% (v/v) ACN, and washed with 0.1% FA before loading the sample and washing again with 0.1% FA. Peptides were eluted from the ZipTips with 0.1% FA in 70% ACN. Finally, samples were dried using a vacuum-centrifuge (Eppendorf) at 30 °C until dryness followed by reconstitution in 12 µL of 0.1% FA in 5% ACN for subsequent MS analysis.

#### LC method for PRM MS

A Thermo Fisher Scientific UltiMate 3000 RSLCnano UHPLC system was used to elute the samples. Each sample was initially injected (6 µL) onto a Thermo Fisher Acclaim PepMap C_18_ trap reversed-phase column (300 µm x 5 mm nano viper, 5 µm particle size) at a flow rate of 15 µL/min using 2% ACN for 3 min with the solvent going to waste. The trap column was switched in-line with the separation column (GRACE Vydac Everest C_18_, 300Å 150 µm x 150 mm, 2 µm) and the peptides were eluted using a flowrate of 3 µL/min using 0.1% FA in water (buffer A) and 80% ACN in buffer A (buffer B) as mobile phases for gradient elution. Following 3 min isocratic of 3% buffer B, peptide elution employed a 3-40% ACN gradient for 28 min followed by 40-95% ACN for 1.5 min and 95% ACN for 1.5 min at 40 °C. The total elution time was 50 min including a 95% ACN wash and a re-equilibration step.

#### PRM MS data acquisition

The eluted peptides from the C18 column were introduced to the MS via a nano-ESI and analysed using the Thermo Fisher Scientific Q-Exactive HF-X mass spectrometer. The electrospray voltage was 1.8 kV in positive ion mode, and the ion transfer tube temperature was 250 °C. Full MS-scans were acquired in the Orbitrap mass analyser over the range m/z 550–560 with a mass resolution of 30,000 (at m/z 200). The AGC target value was set at 1.00E+06 and maximum accumulation time 50 ms for full MS-scans. The PRM inclusion list included two mass values of 552.7640 and 556.7711. MS/MS spectra were acquired in the Orbitrap mass analyser with a mass resolution of 15,000 (at m/z 200). The AGC target value was set at 1.00E+06 and maximum accumulation time 30 ms for MS/MS with an isolation window of 2 m/z. The loop count was set at 14 to gain greater MS/MS data. Raw PRM MS data have been deposited to Panorama at https://panoramaweb.org/Valgepea_Cauto_PRM.url (private reviewer account details: username; password: hSAwNUAL) with a ProteomeXchange Consortium (http://proteomecentral.proteomexchange.org) dataset identifier PXD025760.

#### PRM MS data analysis

Analysis of PRM MS data was performed using the software Skyline^48^. The following parameters were used to extract PRM MS data for the quantification tag sequence GYSFTTTAEK: three precursor isotope peaks with a charge of 2 (++) were included (monoisotopic; M+1; M+2); five of the most intense y product ions from ion 3 to last ion of charge state 1 and 2 among the precursor were picked; chromatograms were extracted with an ion match mass tolerance of 0.05 m/z for product ions by including all matching scans; full trypsin specificity with two missed cleavages allowed for peptides with a length of 8-25 AAs; cysteine carbamidomethylation as a fixed peptide modification. Additionally, peptide modifications included heavy labels for lysine and arginine as ^13^C(6)^15^N(2)/+8.014 Da (K) and ^13^C(6)^15^N(4)/+10.008 Da (R), respectively. This translated into the SIL-proteins and the non-labelled AQUA^®^ peptide possessing the tag GYSFTTTAEK with m/z of 556.7711 and 552.7640, respectively. Hence, the concentrations of SIL-protein stocks were calculated based on the ratio of heavy-to-light GYSFTTTAEK signals and the spike-in of 2 pmol of the non-labelled AQUA^®^ peptide (see above). High accuracy of quantification was evidenced by the very high similarity between both precursor peak areas and expected isotope distribution (R^2^>0.99; idotp in Skyline) and heavy and light peak areas (R^2^>0.99; rdotp in Skyline) for all SIL-protein standard stocks. No heavy GYSFTTTAEK signal was detected for protein CAETHG_RS16140 (an acetylating acetaldehyde dehydrogenase in the NCBI annotation of sequence NC_022592.1^67^), thus 19 SIL-proteins could be used for following absolute proteome quantification in *C. autoethanogenum* (Supplementary Table 1). PRM MS data with all Skyline processing settings can be viewed and downloaded from Panorama at https://panoramaweb.org/Valgepea_Cauto_PRM.url (private reviewer account details: username; password: hSAwNUAL).

### Absolute proteome quantification in *C. autoethanogenum* using DIA MS

We used 19 synthetic heavy SIL variants (see above) of key *C. autoethanogenum* proteins (Supplementary Table 1) as spike-in standards for quantification of intracellular concentrations of their non-SIL counterparts using a DIA MS approach^46^. Also, we performed a dilution series experiment for the spike-in SIL-proteins to ensure accurate absolute quantification. We refer to these 19 intracellular proteins as anchor proteins that were further used to estimate proteome-wide absolute protein concentrations in *C. autoethanogenum*. This was achieved by determining the best linear fit between anchor protein concentrations and their measured DIA MS intensities using the same strategy as described previously^28^.

#### Preparation of spike-in SIL-protein standard mix and dilution series samples

Only Eppendorf Protein LoBind^®^ tubes and pipette tips were used for all preparation steps. The 19 spike-in SIL-protein standards that could be used for absolute proteome quantification in *C. autoethanogenum* (see above) were mixed in two lots: 1) ‘sample spike-in standard mix’: SIL-protein quantities matching estimated intracellular anchor protein quantities (i.e., expected light-to-heavy [L/H] ratios of ∼1) based on label-free absolute quantification of the same samples in our previous work^18^; 2) ‘dilution series standard mix’: SIL-protein quantities doubling the estimated intracellular anchor protein quantities for the dilution series sample with the highest SIL-protein concentrations.

To ensure accurate absolute quantification of anchor protein concentrations, a dilution series experiment was performed to determine the linear dynamic quantification range and LLOQ for each of the 19 spike-in SIL-proteins. Dilution series samples were prepared by making nine 2-fold dilutions of the ‘dilution series spike-in standard mix’ (i.e., 10 samples total for dilution series with a 512-fold concentration span) in a constant *C. autoethanogenum* cell lysate background (0.07 µg/µL; 10 µg/tube) serving as a blocking agent to avoid loss of purified SIL-proteins (to container and pipette tip walls) and as a background proteome for accurate MS quantification of the linear range and LLOQ for anchor proteins.

#### Sample preparation

*C. autoethanogenum* cultures were sampled for proteomics by immediate pelleting of 2 mL of culture using centrifugation (25,000 × *g* for 1 min at 4 °C) and stored at -80 °C until analysis. Details of protein extraction and protein quantification in cell lysates are described previously^18^. In short, thawed cell pellets were suspended in lysis buffer (containing SDS, DTT, and Trizma^®^ base) and cell lysis was performed using glass beads and repeating a ‘lysis cycle’ consisting of heating, bead beating, centrifugation, and vortexing before protein quantification using the 2D Quant Kit (GE Healthcare Life Sciences).

Sample preparation and protein digestion for MS analysis was based on the filter-aided sample preparation (FASP) protocol^68^. The following starting material was loaded onto an Amicon^®^ Ultra-0.5 mL centrifugal filter unit (nominal molecular weight cut-off of 30,000; Merck Millipore): 1) 50 µg of protein for one culture sample from each gas mixture (CO, syngas, or high-H_2_ CO) for building the spectral library for DIA MS data analysis (samples 1-3); 2) 7 µg of protein for one culture sample from either syngas or high-H_2_ CO plus ‘sample spike-in standard mix’ for including spike-in SIL-protein data to the spectral library (samples 4-5); 3) 15 µg of protein for all 12 culture samples (biological quadruplicates from CO, syngas, and high-H_2_ CO) plus ‘sample spike-in standard mix’ for performing absolute proteome quantification in *C. autoethanogenum* (samples 6-17); 4) ten dilution series samples with 10-15 µg of total protein (*C. autoethanogenum* cell lysate background plus ‘dilution series spike-in standard mix’) for performing the dilution series experiment for the 19 spike-in SIL-proteins (see above) (samples 18-27).

Samples containing SIL-proteins (samples 4-27) were incubated at 37 °C for 1 h to reduce SIL-protein disulphide bonds (cell lysate contained DTT). Details of the FASP workflow are described before^18^. In short, samples were washed with UA, sulfhydryl groups alkylated with IAA, proteins digested using a Trypsin/Lys-C mix, and peptides eluted from the filter with 60 µL of ammonium bicarbonate. Next, 50 µL of samples 1-3 were withdrawn and pooled for performing high pH reverse-phase fractionation as described previously^18^ for expanding the spectral library for DIA MS data analysis, yielding eight fractions (samples 28-35). Subsequently, all samples were vacuum-centrifuged at 30 °C until dryness followed by reconstitution of samples 1-3 and 4-35 in 51 and 13 µL of 0.1% FA in 5% ACN, respectively. Finally, total peptide concentration in each sample was determined using the Pierce™ Quantitative Fluorometric Peptide Assay (Thermo Fisher Scientific) to ensure that the same total peptide amount across samples 1-17 and 28-35 (excluding samples 18-27, see below) could be injected for DIA MS analysis.

#### LC method for data-dependent acquisition (DDA) and DIA MS

Details of the LC method employed for generating the spectral library using DDA and for DIA sample runs are described previously^20^. In short, a Thermo Fisher Scientific UHPLC system including C_18_ trap and separation columns was used to elute peptides with a gradient and total elution time of 110 min. For each DDA and DIA sample run, 1 µg of peptide material from protein digestion was injected, except for dilution series samples (samples 18-27 above) that were injected in a constant volume of 3 µL to maintain the dilution levels of the ‘dilution series spike-in standard mix’.

#### DDA MS spectral library generation

The following 13 samples were analysed on the Q-Exactive HF-X in DDA mode to yield the spectral library for DIA MS data analysis: 1) three replicates of one culture sample from each gas mixture (CO, syngas, or high-H_2_ CO) (samples 1-3 above); 2) three replicates of one culture sample from either syngas or high-H_2_ CO plus ‘sample spike-in standard mix’ (samples 4-5); 3) eight high pH reverse-phase fractions of a pool of samples from each gas mixture (samples 28-35).

Details of DDA MS acquisition for generating the spectral library are described before^20^. In short, eluted peptides from the C_18_ column were introduced to the MS *via* a nano-ESI and analysed using the Q-Exactive HF-X with an Orbitrap mass analyser. The DDA MS spectral library for DIA MS data confirmation and quantification using the software Skyline^48^ was created using the Proteome Discoverer 2.2 software (Thermo Fisher Scientific) and its SEQUEST HT search as described previously^18^. The final .pd result file contained peptide-spectrum matches (PSMs) with q-values estimated at 1% false discovery rate (FDR) for peptides ≥4 AAs. The generated spectral library file has been deposited to the ProteomeXchange Consortium (http://proteomecentral.proteomexchange.org) *via* the PRIDE partner repository^69^ with the dataset identifier PXD025732 (private reviewer account details: username; password: QdkJHXy0).

#### DIA MS data acquisition

Details of DIA MS acquisition are described before^20^. In short, as for DDA MS acquisition, eluted peptides were introduced to the MS *via* a nano-ESI and analysed using the Q-Exactive HF-X with an Orbitrap mass analyser. DIA was achieved using an inclusion list: m/z 395□1100 in steps of 15 amu and scans cycled through the list of 48 isolation windows with a loop count of 48. In total, DIA MS data was acquired for 22 samples (samples 6-27 defined in section *Sample preparation*). Raw DIA MS data have been deposited to the ProteomeXchange Consortium (http://proteomecentral.proteomexchange.org) *via* the PRIDE partner repository^69^ with the dataset identifier PXD025732 (private reviewer account details: username; password: QdkJHXy0).

#### DIA MS data analysis

DIA MS data analysis was performed with Skyline^48^ as described before^18^ with the following modifications: 1) 12 manually picked high confidence endogenous peptides present in all samples and spanning the elution gradient were used for iRT alignment through building an RT predictor; 2) outlier peptides from iRT regression were removed; 3) a minimum of three isotope peaks were required for a precursor; 4) single peptide per spike-in SIL-protein was allowed for anchor protein absolute quantification while at least two peptides per protein were required for label-free estimation of proteome-wide protein concentrations; 5) extracted ion chromatograms (XICs) were transformed using Savitszky-Golay smoothing. Briefly, the .pd result file from Proteome Discoverer was used to build the DIA MS spectral library and the mProphet peak picking algorithm^47^ within Skyline was used to separate true from false positive peak groups (per sample) and only peak groups with q-value<0.01 (representing 1% FDR) were used for further quantification. We confidently quantitated 7,288 peptides and 1,243 proteins across all samples and 4,887 peptides and 1,043 proteins on average within each sample for estimating proteome-wide absolute protein concentrations. For absolute quantification of anchor protein concentrations, we additionally manually: 1) removed integration of peaks showing non-Gaussian shapes or interference from other peaks; 2) removed precursors with similarity measures of R^2^<0.9 between product peak areas and corresponding intensities in the spectral library (dotp in Skyline), precursor peak areas and expected isotope distribution (idotp), or heavy and light peak areas (rdotp). After analysis in Skyline, 17 spike-in SIL-proteins remained for further analysis as protein CAETHG_RS14410 was not identified in DIA MS data while CAETHG_RS18395 did not pass quantification filters (Supplementary Table 1). DIA MS data with all Skyline processing settings can be viewed and downloaded from Panorama at https://panoramaweb.org/Valgepea_Cauto_Anchors.url for anchor protein absolute quantification and at https://panoramaweb.org/Valgepea_Cauto_LF.url for estimating proteome-wide absolute protein concentrations (private reviewer account details: username; password: hSAwNUAL).

#### Absolute quantification of anchor protein concentrations

We employed further stringent criteria on top of the output from Skyline analysis to ensure high confidence absolute quantification of 17 anchor protein concentrations. Firstly, precursor with highest heavy intensity for the highest ‘dilution series spike-in standard mix’ sample in the dilution series (DS01) was kept while others were deleted. Peptides quantified in less than three biological replicates cultures within a gas mixture, with no heavy signal for DS01 sample, or with heavy signals for less than three continuous dilution series samples were removed. Next, we utilised the dilution series experiment to only keep signals over the LLOQ and within the linear dynamic quantification range. For this, correlation between experimental and expected peptide L/H signal ratios for each peptide across the dilution series was made to determine the LLOQ and calculate correlation, slope, and intercept between MS signal and spike-in level (Supplementary Table 2). Only peptides showing correlation R^2^>0.95, 0.95<slope<1.05, and -0.1<intercept<0.1 for the dilution series were kept. This ensured that we were only using peptides within the linear dynamic range. The remaining peptides were further filtered for each culture sample by removing peptides whose light or heavy signal was below the LLOQ in the dilution series. Subsequently, only peptides were kept with L/H ratios for at least three biological replicates cultures for each gas mixture (i.e., ≥9 data points). Finally, we aimed to detect outlier peptides by calculating the % difference of a peptide’s L/H ratio from the average L/H ratio of all peptides for a given protein for every sample. Peptides were considered outliers and thus removed if the average difference across all samples was >50% or if the average difference within biological replicate cultures was >50%. After the previous stringent criteria were applied, 106 high-confidence peptides remained (Supplementary Table 2) for the quantification of 16 anchor protein concentrations since CAETHG_RS01830 was lost during manual analysis (Table 1 and Supplementary Table 1). Proteins CAETHG_RS13725 and CAETHG_RS07840 were excluded from the high-H_2_ CO culture dataset as their calculated concentrations varied >50% between biological replicates. Data of one high-H_2_ CO culture was excluded from further analysis due to difference from bio-replicates likely due to challenges with MS analysis.

#### Label-free estimation of proteome-wide protein concentrations

We used the anchor proteins to estimate proteome-wide protein concentrations in *C. autoethanogenum* by determining the best linear fit between anchor protein concentrations and their measured DIA MS intensities using the aLFQ R package^49^ and the same strategy as described for SWATH MS^28^. Briefly, aLFQ used anchor proteins and cross-validated model selection by bootstrapping to determine the optimal model within various label-free absolute proteome quantification approaches (e.g., TopN, iBAQ). The approach can obtain the model with the smallest error between anchor protein concentrations determined using SIL-protein standards and label-free estimated concentrations. The models with the highest accuracy were used to estimate proteome-wide label-free concentrations for all proteins from their DIA MS intensities (1,043 proteins on average within each sample; minimal two peptides per protein; see above): summing the five most intense fragment ion intensities of the most or three of the most intense peptides per protein for CO or high-H_2_ CO cultures, respectively; summing the five most intense fragment ion intensities of all quantified peptides of the protein divided by the number of theoretically observable peptides (i.e., iBAQ^70^) for syngas cultures.

### Expected protein complex stoichiometries

Equimolar stoichiometries for the HytA-E/FdhA and MetFV protein complexes were expected based on SDS gel staining experiments in *C. autoethanogenum*^57^ and the acetogen *Moorella thermoacetica*^71^, respectively. Expected stoichiometries for other protein complexes in Fig. 3 and Supplementary Fig. 3 were based on measured stoichiometries in *E. coli* K-12 (Complex Portal; www.ebi.ac.uk/complexportal) and significant homology between complex member proteins in *C. autoethanogenum* and *E. coli*. All depicted *C. autoethanogenum* protein complex members had NCBI protein-protein BLAST E-values<10^−16^ and scores>73 against respective *E. coli* K-12 proteins using non-redundant protein sequences.

### Generation of proteomaps

The distribution of proteome-wide protein concentrations among functional gene classifications was visualised using proteomaps^27^. For this, the NCBI annotation of sequence NC_022592.1^67^ was used as the annotation genome for *C. autoethanogenum*, with CAETHG_RS07860 removed and replaced with the carbon monoxide dehydrogenase genes named CAETHG_RS07861 and CAETHG_RS07862 with initial IDs of CAETHG_1620 and 1621, respectively. Functional categories were assigned to protein sequences with KO IDs^51^ using the KEGG annotation tool BlastKOALA^72^. Since proteomaps require a tree-like hierarchy, proteins that were automatically assigned to multiple functional categories were manually assigned to one bottom-level category (Level 3 in Supplementary Table 4) based on their principal task. We also created functional categories “C1 fixation/Wood-Ljungdahl Pathway” (Level 2/3), “Acetate & ethanol synthesis” (Level 3), “Energy conservation” (Level 3), “Hydrogenases” (Level 3) and manually assigned key acetogen proteins to these categories to reflect more accurately functional categories for an acetogen. Proteins without designated KO IDs were manually assigned to latterly created categories or grouped under “Not Included in Pathway or Brite” (Level 1) with Level 2 and 3 as “No KO ID”. If BlastKOALA assigned multiple genes the same proposed gene/protein name, unique numbers were added to names (e.g., pfkA, pfkA2). The final “treemap” defining the hierarchy for our proteomaps is in Supplementary Table 4.

### Calculation of apparent in vivo catalytic rates of enzymes (k_app_)

We calculated apparent in vivo catalytic rates of enzymes, denoted as k_app_ (s^-1^)^26^, as the ratio of specific flux rate (mmol/gDCW/h) determined before^18^ and protein concentration (nmol/gDCW) quantified here for the same *C. autoethanogenum* CO, syngas, and high-H_2_ CO cultures. Gene-protein-reaction (GPR) data of the genome-scale metabolic model iCLAU786^18^ were manually curated to reflect most recent knowledge and were used to link metabolic fluxes with catalysing enzymes. For reactions with multiple assigned enzymes (i.e,. isoenzymes), the enzyme with the highest average ranking of its concentration across the three cultures (Supplementary Table 3) was assumed to solely catalyse the flux. For enzyme complexes, average of quantified subunit concentrations was used (standard deviation estimated using error propagation). For the HytA-E hydrogenase, its measured protein concentration was split between reactions “rxn08518_c0” (direct CO_2_ reduction with H_2_ in complex with FdhA) and “leq000001_c0” (H_2_ oxidation solely by HytA-E) proportionally to flux for syngas and high-H_2_ CO cultures. The resulting enzymes or enzyme complexes catalysing specific fluxes are shown in Supplementary Table 5. Finally, we assumed each protein chain being catalytically active and only calculated k_app_ values for metabolic reactions with a non-zero flux in at least one condition, specific flux rate>0.1% of CO fixation flux in at least one condition, and label-free data with measured concentration for the associated enzyme(s) in all conditions (Supplementary Table 5). Membrane proteins were excluded from k_app_ calculations to avoid bias from potentially incomplete protein extraction. This produced k_app_ values for 13 enzymes/complexes using anchor protein concentrations and for 48 enzymes/complexes using label-free protein concentrations (Supplementary Table 5).

### Determination of regulation level of metabolic fluxes

We used published flux and relative proteomics data^18^ of the same cultures studied here to determine whether fluxes are regulated by changing enzyme concentrations or their catalytic rates by considering metabolic fluxes with non-zero specific flux rates in at least two conditions of CO, syngas, or high-H_2_ CO cultures. The same manually curated GPRs and criteria for isoenzymes and protein complexes as described above for k_app_ calculation were used to determine flux-enzyme pairs (Supplementary Table 6). We first used a two-tailed two-sample equal variance Student’s t-test with FDR correction^73^ to determine fluxes with

significant changes between any two conditions (q-value<0.05). We then used the Student’s left-tailed t-distribution with FDR to determine if the significant flux change for every flux was significantly different from the change in respective enzyme expression between the same conditions (Supplementary Table 6). Flux with a q-value<0.05 for the latter test was considered to be regulated at post-translational level (e.g., by changing enzyme catalytic rate).

## DATA AVAILABILITY

All data generated or analysed during this study are in the main text, supplementary information files, or public databases. Raw PRM MS data have been deposited to Panorama at https://panoramaweb.org/Valgepea_Cauto_PRM.url (private reviewer account details: username; password: hSAwNUAL) with a ProteomeXchange Consortium (http://proteomecentral.proteomexchange.org) dataset identifier PXD025760. Raw DIA MS data have been deposited to the ProteomeXchange Consortium (http://proteomecentral.proteomexchange.org) *via* the PRIDE partner repository^69^ with the dataset identifier PXD025732 (private reviewer account details: username; password: QdkJHXy0). PRM MS data with all Skyline processing settings can be viewed and downloaded from Panorama at https://panoramaweb.org/Valgepea_Cauto_PRM.url. DIA MS data with all Skyline processing settings can be viewed and downloaded from Panorama at https://panoramaweb.org/Valgepea_Cauto_Anchors.url for anchor protein absolute quantification and at https://panoramaweb.org/Valgepea_Cauto_LF.url for estimating proteome-wide absolute protein concentrations (private reviewer account details: username; password: hSAwNUAL). Any other relevant data are available from the corresponding author on reasonable request.

## ACKNOWLEDGEMENTS

We thank Jörg Bernhardt for help with proteomaps, Tim McCubbin for scripts, Andrus Seiman for statistics, and Olivier Lemaire for valuable discussions. This work was funded by the Australian Research Council (ARC LP140100213) in collaboration with LanzaTech. We thank the following investors in LanzaTech’s technology: Sir Stephen Tindall, Khosla Ventures, Qiming Venture Partners, Softbank China, the Malaysian Life Sciences Capital Fund, Mitsui, Primetals, CICC Growth Capital Fund I, L.P. and the New Zealand Superannuation Fund. The research utilised equipment and support provided by the Queensland node of Metabolomics Australia, an initiative of the Australian Government being conducted as part of the NCRIS National Research Infrastructure for Australia. There was no funding support from the European Union for the experimental part of the study. However, K.V. acknowledges support also from the European Union’s Horizon 2020 research and innovation programme under grant agreement N810755.

## AUTHOR CONTRIBUTIONS

Conceptualisation, K.V., C.L., M.K., L.K.N., and E.M.; Methodology, K.V., G.T., N.T., A.T., C.L., A.P.M., R.T., and E.M.; Formal Analysis, K.V., and G.T.; Investigation, K.V., G.T., N.T., and A.T.; Resources, N.T., A.T., M.K., S.D.S., L.K.N., and E.M.; Writing – Original Draft, K.V. and E.M.; Writing – Review & Editing, K.V., N.T., C.L., A.P.M., R.T., M.K., L.K.N., and E.M.; Supervision, M.K., L.K.N., and E.M.; Project Administration, R.T., and E.M.; Funding Acquisition, M.K., S.D.S., L.K.N., and E.M..

